# Reconciling persistent and dynamic hypotheses of working memory coding in prefrontal cortex

**DOI:** 10.1101/231506

**Authors:** SE Cavanagh, JP Towers, JD Wallis, LT Hunt, SW Kennerley

## Abstract

Competing accounts propose that working memory (WM) is subserved either by persistent activity in single neurons or by dynamic (time-varying) activity across a neural population. Here we compare these hypotheses across four regions of prefrontal cortex (PFC) in a spatial WM task, where an intervening distractor indicated the reward available for a correct saccade. WM representations were strongest in ventrolateral PFC (VLPFC) neurons with higher intrinsic temporal stability (time-constant). At the population-level, although a stable mnemonic state was reached during the delay, this tuning geometry was reversed relative to cue-period selectivity, and was disrupted by the distractor. Single-neuron analysis revealed many neurons switched to coding reward, rather than maintaining task-relevant spatial selectivity until saccade. These results imply WM is fulfilled by dynamic, population-level activity within high time-constant neurons. Rather than persistent activity supporting stable mnemonic representations that bridge distraction, PFC neurons may stabilise a dynamic population-level process that supports WM.

---

Temporary maintenance of relevant information in the absence of direct sensory input is a crucial component of working memory (WM). The neuronal basis of WM has been studied extensively through single-neuron recordings. These typically involve non-human primates performing tasks where a transient sensory stimulus must be remembered across a several second delay before a probe cues a response to the remembered stimulus. A consensus has developed from these experiments^1-3^, and from lesion studies^4,5^, that cognitive operations that use information in WM depend upon the prefrontal cortex (PFC) ^6^, with individual PFC neurons sustaining stimulus-specific representations across the mnemonic delay. This *stable coding* has inspired biophysically plausible attractor network models of working memory, in which persistent activity is facilitated by a neocortical circuit structured with strong recurrent connections between similarly tuned neurons^7^.

Recent findings have challenged these established views. Responses of PFC neurons are often highly heterogeneous, with only a minority exhibiting prolonged stimulus-specific encoding during a delay^8-12^. The majority of neurons instead show short-lived selectivity, with variable onset latencies and durations. This pattern of working memory activity is referred to as *dynamic coding*. Evidence for dynamic coding has led to revised attractor models that reconcile time-varying and stable single neuron responses^13^. It has also inspired alternate explanations for how WM may be achieved without relying upon a stable representation in the form of persistent spiking activity^8,14-18^. These include *dynamic trajectory* models where neural firing preserves a representation of the mnemonic stimulus throughout a delay by moving through a reproducible path of activity^15,17,18^. They also include *synaptic* models where WM is achieved by short-term plasticity of synaptic weights^8,14^. In the latter, stable delay-period WM correlates still arise, but as a by-product of spontaneous activity within a circuit that is temporarily embedded with mnemonic information.

An important prediction rarely tested in the context of WM models relates to how network representations of stimuli resist distraction^19-21^. In a world where we are constantly exposed to salient sensory stimuli, efficient cognitive operations that depend on WM require that this information is resistant to distractions in our environment. The majority of task designs used to study single-neuron WM-correlates lack intervening stimuli during delays. If memoranda are maintained purely by persistent single neuron activity, and if those neurons flexibly encode multiple task features (as is common in PFC^22-27^), a distracting stimulus could disrupt the attractor state and cause the memory to be distorted or lost. Several neurophysiological accounts suggest PFC possesses a privileged position in cortical processing – the ability for individual task-selective neurons to resist distraction^28-30^. More recently, however, the view that PFC neurons are resistant to distractors has been challenged^21,31^. If WM is maintained in the absence of stable single-neuron representations, it becomes important to understand how memoranda are encoded across the PFC population in the face of distraction, and what role neurons with persistent activity play in such population-level encoding.

One factor worth considering is that single neurons exhibit considerable heterogeneity in the degree to which they exhibit persistent activity at rest^32,33^. By fitting an exponential decay to the autocorrelation of neuronal firing outside of the task, it is possible to characterise individual neurons’ intrinsic temporal stability (time constant)^33,34^. A neuron’s time constant likely reflects a combination of its intrinsic physical properties and its degree of recurrent connectivity^35^. Because neurons with higher time constants were more likely to be maintain information during extended cognitive processes such as decision-making^33^, we hypothesised that heterogeneity in single-neuron time constants may explain why some neurons retain stimulus-specific mnemonic representations over a delay, whereas others exhibit more transient selectivity. This would reconcile persistent and dynamic WM coding. If attractor states underlie WM, classical stable mnemonic representations should primarily be evident in a subpopulation of neurons with high time constants. Furthermore, neurons with high time constants may facilitate the stability of WM representations throughout distraction.

We tested these hypotheses in a spatial WM task where a stimulus revealing the reward for a correct response was presented either before or after the spatial WM cue. Presentation after the mnemonic cue serves as a salient distractor, potentially disrupting spatial WM representations^36,37^. This also allowed us to test how an interfering stimulus affected network-level mnemonic coding as a function of neuronal time constant.

## Results

### Task and Neurophysiological Recordings

Two rhesus macaques *(Macaca mulatta)* performed a spatial working memory task where the reward amount for successful responses varied across trials (**Fig1a**)^36,37^. Briefly (see **Methods**), subjects were first required to fixate a central cue for 1000ms. If fixation was maintained, two cues were sequentially presented (for 500ms apiece), each followed by a 1000ms delay. The spatial cue indicated which of 24 locations the subject had to hold in working memory (the mnemonic stimulus); the reward cue indicated which of 5 reward magnitudes the subject would receive for a saccade to the remembered location. The subject could elicit a saccade to the remembered location following a go cue. In “RS trials”, the first and second cues were the reward and spatial cues respectively; the cue order was reversed in “SR trials”. We counterbalanced all spatial positions and reward levels, and the two trial types were randomly intermingled. As only the spatial cue was relevant for correct performance, the reward cue on SR trials may serve as a distracting stimulus, interfering with the stability of spatial working memory representations.

**Figure 1:**
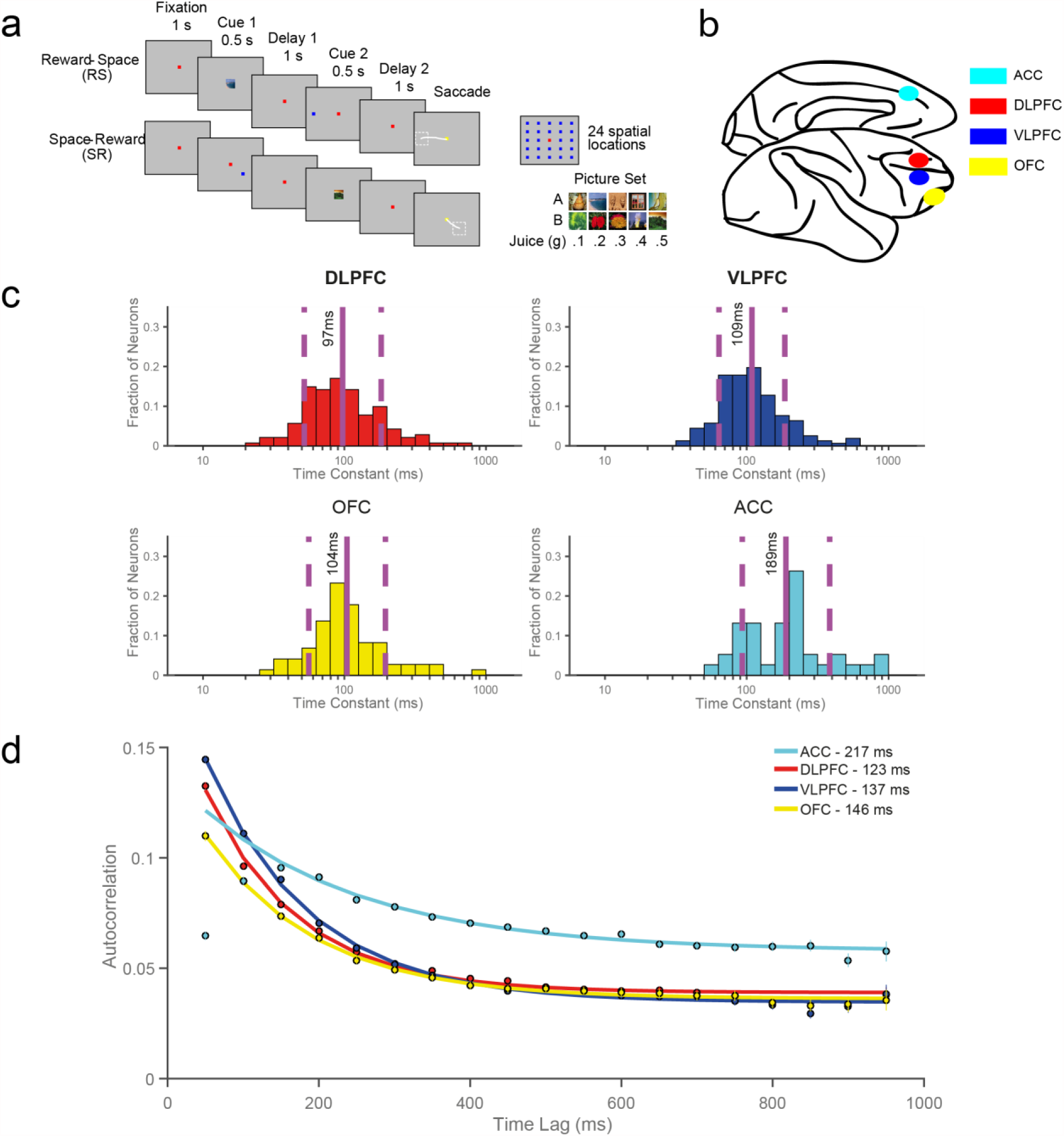
Overview of reward-varying spatial working-memory task, recording locations and time constant analysis. **a)** Reward-varying spatial working memory task. Monkeys were trained to remember a spatial position in working memory. They were also presented with a cue indicating the reward size they would receive for successfully completing the trial with a saccade to the remembered location. On RS (Reward-Space) trials, the reward cue was presented first; whereas on the SR (Space-Reward) trials, the cues were presented in the reverse order. On SR trials the reward cue therefore acted as a distraction to working memory of the task-relevant spatial information. **b)** Approximate location of neural recordings. Neurons were recorded from anterior cingulate cortex (ACC), dorsolateral prefrontal cortex (DLPFC), ventrolateral prefrontal cortex (VLPFC), and orbitofrontal cortex (OFC). **c)** Histograms of the single-neuron time constants within the four PFC brain regions. Time constants are highly variable across neurons. Solid and dashed vertical lines represent mean(Log(τ)) and mean(Log(τ)) ± SD(Log(τ)) respectively. **d)** Population-level time constants of firing rate autocorrelation in DLPFC, VLPFC, OFC and ACC during pre-stimulus fixation epoch. Time constant captures the rate of decay of autocorrelation over time. ACC had the highest and most distinct time constant of all PFC regions studied.

Single neurons were recorded from four brain regions across prefrontal cortex (PFC; **Fig1b**): anterior cingulate cortex (ACC; areas 9m, 24c, n= 198), dorsolateral PFC (DLPFC; areas 9, 9/46d, n= 209), ventrolateral PFC (VLPFC; areas 9/46v, 45A, n= 206) and orbitofrontal cortex (OFC; areas 11, 13, n= 152). Histological reconstruction of recording locations is reported elsewhere^36,37^. All neurons per region were pooled across sessions to form “pseudopopulations” in order to examine population-level activity^12,13,38^. Importantly, neurons were not pre-screened for functional properties prior to recordings, facilitating an unbiased examination of population coding-dynamics and single-neuron resting time constant measures.

### Resting time constants

We first sought to define each neuron’s resting time constant (“tau”) by fitting an exponential decay to its spike-count autocorrelation during the 1000ms fixation period^33^. The autocorrelation functions of those neurons that could successfully be described by an exponential decay with an offset^34^ were fitted to yield a resting time constant for each neuron (409 of 765 single neurons, see **Methods**).

As previously reported^33^, there was marked heterogeneity in the temporal specialisation of individual neurons both within and between PFC regions (**Fig1c**). Time constants differed significantly across areas (Kruskal-Wallis test, p=2.21 x 10^−6^), where the highest taus were within the ACC population (Mann-Witney U Tests; ACC v DLPFC, 4.51 x 10^−7^; ACC v OFC, 2.48 x 10^−5^; ACC v VLPFC, 7.58 x 10^−6^). We next characterised the population-level taus of the four PFC brain regions (**Fig1d**, see **Methods**). For this analysis, data from all recorded neurons within each brain area was fitted using the same exponential decay equation. This approach has previously shown a hierarchy of temporal specialisations exists across cortex^34^. Our results were consistent with this, again emphasising the distinction of ACC at the summit of a hierarchy across PFC regions^33,34^.

### Decoding analysis of Working Memory Activity

We next applied a multivariate decoding approach to investigate population-dynamics across PFC^38^. Briefly, this involved calculating the average single-neuron firing rate for each condition (8 collapsed locations for Space; 5 Reward levels; see **Methods**) within two independent halves of the data (training and test sets). The difference in mean activity between each pair of conditions was calculated within each set (e.g. 28 pairwise differences for Space, 10 pairwise differences for Reward). For all neurons within each regional pseudopopulation, each pairwise conditional difference was correlated between the training and test sets to quantify how well each PFC region’s activity discriminated between the different conditions.

Our results provide the most complete comparison to date of population-level WM activity patterns across multiple PFC brain regions (**Fig2**). Of the four PFC regions examined, VLPFC activity best discriminated between both the different spatial locations and the different reward sizes regardless of trial (SR, RS) type, and it was the only PFC region that sustained these selectivity patterns across delays. VLPFC was also the PFC region most strongly discriminating spatial information immediately prior to saccade. However, VLPFC activity exhibited a distinct temporal profile. On the SR task, spatial information was strongly represented during both the spatial cue and the first delay (**Fig2a**; Spatial cue and Delay One, p <0.0001; cluster-based permutation tests). Importantly, shortly after the reward cue was presented in SR trials, the VLPFC spatial discriminability was dramatically reduced (**Fig2a**, Delay Two). Instead, a robust representation of reward emerged which was maintained through to the end of the trial (**Fig2b**; Reward cue through end of trial, p <0.0001; cluster-based permutation tests). This strong reward representation, seemingly at the expense of the spatial WM representation, was noteworthy, as retaining a memory of the spatial location is the key task variable necessary for correct performance. A similar pattern of selectivity switching was present in RS trials, where the VLPFC population initially maintained a representation of the expected reward, but this representation attenuated as the spatial representation strongly emerged following the spatial cue (**Fig2c**, Spatial cue and Delay Two, p <0.0001; **Fig2d**, Reward cue and Delay One, p <0.0001; cluster-based permutation tests). These results are consistent with VLPFC spiking-activity prioritising a representation of the most recently attended information, regardless of whether it is necessary to store the stimulus in working memory for successful performance^39,40^.

**Figure 2:**
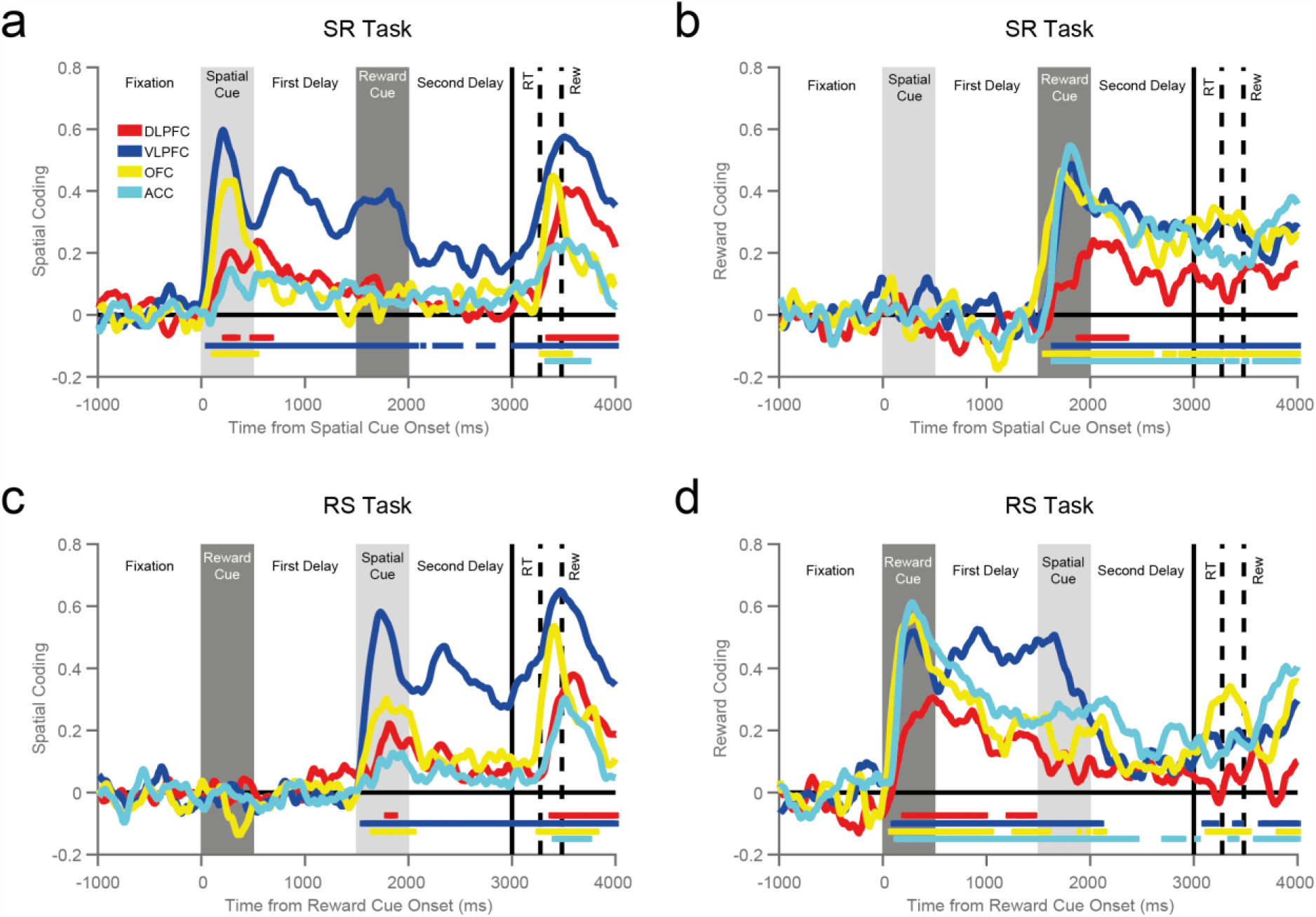
Ventrolateral prefrontal neurons maintain information for both spatial and reward stimuli during delay epochs. The coding of each task feature (spatial location, **a** and **c**; reward level, **b** and **d**) are plotted for each brain area and trial type (SR-task, **a** and **b**; RS-task, **c** and **d**). Ventrolateral prefrontal cortex (VLPFC) is the only region to strongly code information about space and reward across the trial. Notably, the VLPFC activity primarily encodes the factor most recently presented. When the reward cue is shown first (RS task, **c** and **d**), a representation of reward size is maintained throughout the first delay, but falls away when the spatial cue is presented. More surprisingly, a similar weakening of spatial coding is also observed on the SR Task (**a**), even though this analysis is restricted to trials where the subject remembered the correct spatial location. Therefore, the maintenance of a strong population code for spatial location within this epoch does not seem essential for working memory. The VLPFC population strongly encodes and maintains a representation of the remembered spatial location, but this is substantially weakened by the offset of the reward cue. The first solid vertical line signifies when subjects were cued to respond. The first and second dashed vertical lines represent the average timing of the subjects’ saccade and the onset of reward respectively. Coloured horizontal lines represent significant encoding for the corresponding brain region (Cluster-based permutation test, p<0.05).

Maintenance of spatial discriminability in DLPFC was weak, emerging relatively late in the spatial cue epoch and decaying shortly after the first delay (**Fig2a, c**). This is surprising given that DLPFC has often been implicated in the stable maintenance of working memory, but such discrepancies may be due to variability in recordings along the anterior-posterior gradient of DLPFC^41^, or studies describing DLPFC cells or lesions which extend to surrounding areas including VLPFC^1,4,5^. OFC had phasic representations of spatial location during cue presentation and response^42^ (**Fig2a, c**), while ACC only exhibited brief spatial selectivity at the time of reward. ACC, OFC and VLPFC all had prolonged maintenance of reward size in both trial types (**Fig2b, d**), consistent with ACC and OFC playing a key role in reward-guided behaviour^33,43-45^.

### Population-activity separated by resting time constant

We next sought to link the two previous analyses, exploring whether the heterogeneity of single-neuron time constants (**Fig1c**) predicted different functional roles during working memory. As cells with higher taus have an intrinsic capacity for sustained persistent activity, we hypothesised that these cells would more likely be integral to stable attractor states and therefore exhibit stronger and more prolonged maintenance of spatial information^7,13^. We focussed upon VLPFC, as this was the only candidate region with sustained spatial selectivity. We subdivided the population based upon a median split of tau^33^, and then re-computed the spatial and reward discriminability as in **Fig2** for high and low tau subpopulations (**Fig3**).

**Figure 3:**
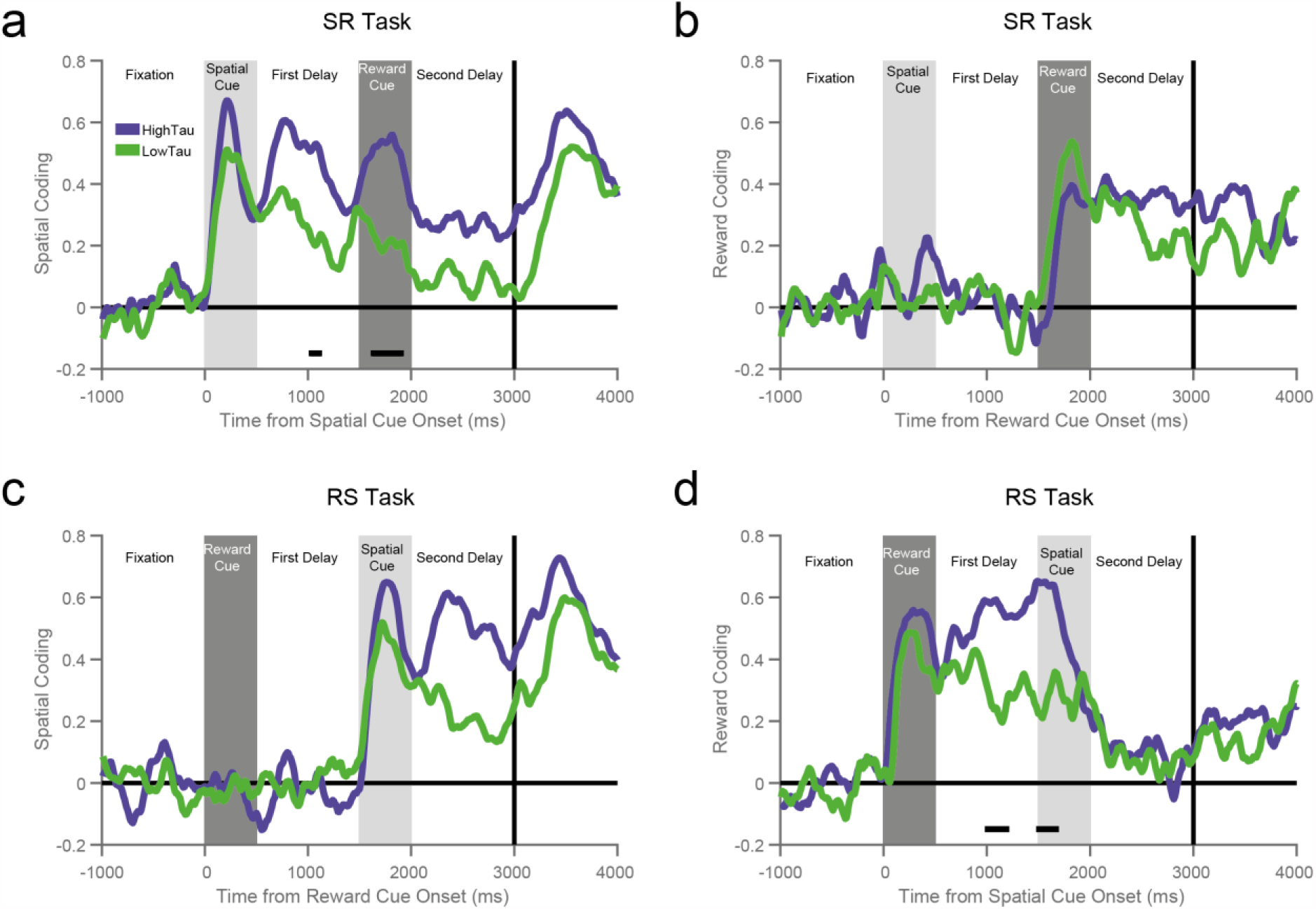
Ventrolateral prefrontal neurons with higher resting time constants maintain reward and spatial information across delays. Coding for space (**a** and **c**) and reward size (**b** and **d**) is calculated for two subpopulations of ventrolateral prefrontal cortex (VLPFC) neurons subdivided by resting time constant. The subpopulation with higher time constants has a stronger representation of remembered spatial location during the first delay of the SR task (**a**, p = 0.0482) and whilst the reward cue is on screen (p = 0.0027). The high time constant population also has a trend towards having stronger spatial coding in the second delay of the RS task (**c**, p = 0.0639). These neurons code reward more strongly during the first delay of the RS task (**d**, p = 0.0457) and just as the spatial cue is being presented (p = 0.0077). Notably, on SR trials, where the reward cue is acting as a distractor, the high time constant subpopulation do not exhibit stronger reward coding. They also switch off reward coding on RS trials as soon as the task-relevant (spatial) cue is presented. Horizontal black bars represent a significant difference between the high and low time constant subpopulations (Cluster-based permutation test, p<0.05).

As hypothesised, the high tau VLPFC neuronal subpopulation had more sustained selectivity for both spatial and reward information. Both tau subpopulations showed a similar temporal profile to the whole population of VLPFC neurons, but selectivity in the low tau population decayed quickly following stimulus offset. A formal comparison between the two subpopulations indicated the high tau subpopulation had stronger spatial selectivity during delay one (p=0.0482, cluster based permutation test) and reward cue presentation (p = 0.0027) of the SR task (**Fig3a**), and stronger reward coding during delay one (p = 0.0457) and when the spatial cue was presented (p=0.0077) on the RS task (**Fig3d**). However, an examination of activity during the task epoch when the respective reward or spatial cue was onscreen revealed strong selectivity that was statistically indistinguishable between the two subpopulations (“spatial cue” of **Fig3a,c**; “reward cue” of **Fig3b,d**). In other words, it is not the case that low tau subpopulations are simply less task-selective. Instead, high tau cells appear to be specialised for exhibiting sustained selectivity across delays, a property which may be critical for supporting WM processes.

### Cross-temporal activity separated by resting time constant

The results presented so far – sustained population-level selectivity across delays only in cells with persistent resting activity - could be explained by both attractor models and alternate hypotheses of working memory^7,46^. They are also consistent with previous reports relating baseline autocorrelation to WM activity in single neurons^47^. The population WM selectivity in **Fig3** could be supported either by individual neurons maintaining strong selectivity across the trial, or neurons dynamically encoding information with different latencies and durations such that the population-level selectivity is maintained over time.

To contrast between these hypotheses, we performed a cross-temporal pattern analysis to probe the stability of the active encoding state^33,38^. To study cross-temporal generalisation of task selectivity, a classifier is trained at one timepoint (*t*) and tested at a different timepoint (*t+ δ*). If there remains a strong correlation between the test and training set at two distinct timepoints, selectivity generalises across the period between the two timepoints. By using all ***n*** timepoints as training and test sets, an ***n × n*** correlation matrix can be constructed.

**Figure 4:**
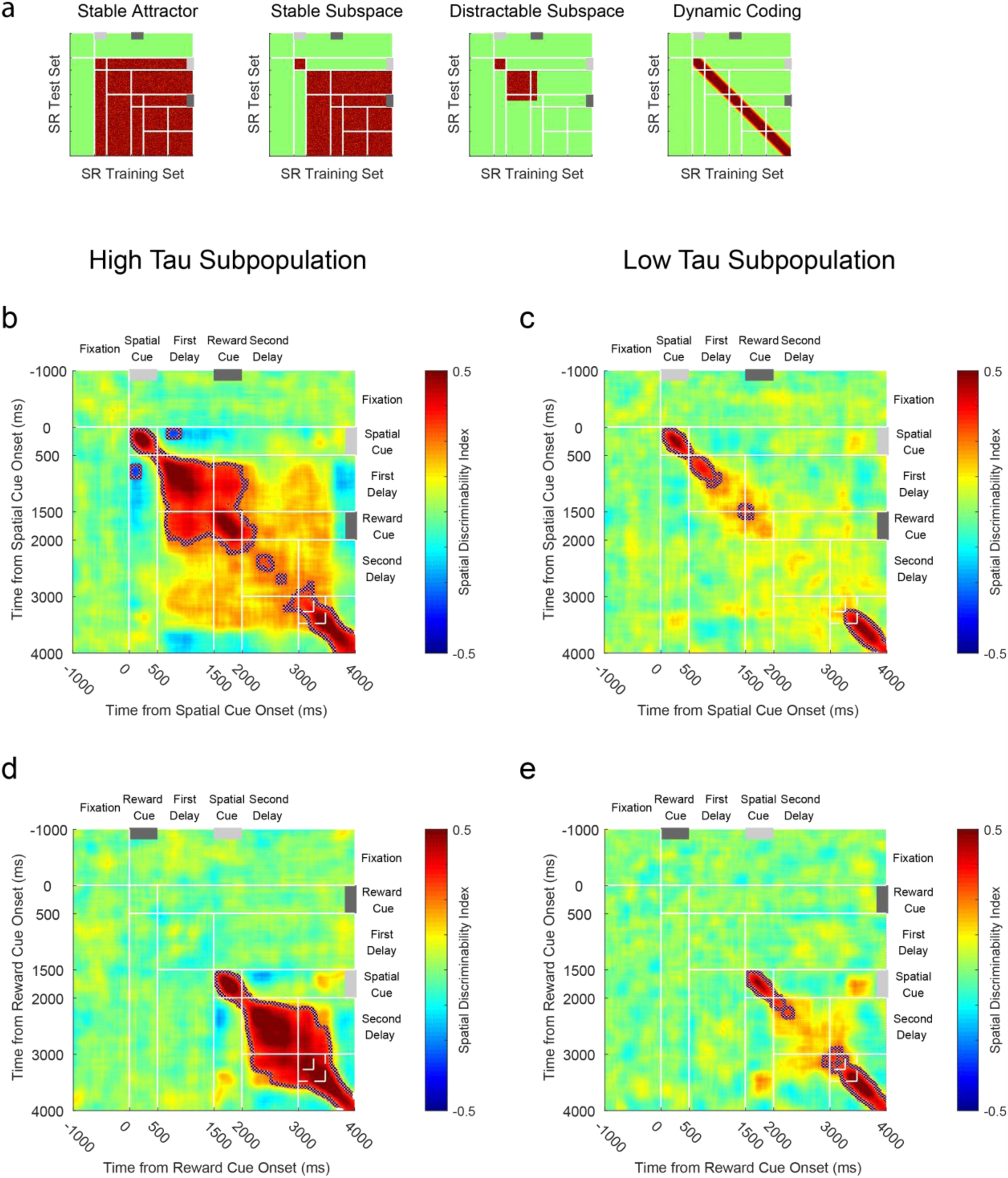
Cross-temporal dynamics of spatial selectivity by high and low time constant populations. **a)** Schematic representing cross-temporal dynamics of different working-memory codes on SR trials. Each pixel represents how well spatial location can be discriminated when using half of the trials at one timepoint as a training-set (X-Axis), and the other half of trials at a separate timepoint as a test-set (Y-Axis). On diagonal, the value is identical to those plotted in Fig3. Off diagonal, the plot indicates the stability of any spatial coding across time. In the first exemplar, stable spatial coding is evident across the trial, as data from any timepoint after cue presentation can be used to decode the remembered spatial location at any other timepoint. The second exemplar is similar, but this stable state is only established following a transient dynamic phase where the cue is initially encoded. The third exemplar shows that this stable state is established during the initial delay – but collapses after the reward cue is presented. The final exemplar shows that spatial location is coded throughout the trial (heat on the diagonal), but that this code is not stable across time. Therefore, the way space is coded at two distinct timepoints is inconsistent. **b-e)** Cross-temporal decodability of spatial location is plotted for high (**b, d**) and low (**c, e**) time constant VLPFC populations on SR (**b, c**) and RS **(d, e)** trials. The high time constant subpopulation has a much greater stability of its spatial coding: the off-diagonal elements are warm, meaning that the same population code persists throughout the delay epoch following the spatial cue. Despite this stability, there is a negative correlation between the cue period and the delay indicating a reversal of spatial tuning between these epochs. In SR trials, a stable state is reached during the first delay, but this is disrupted by the presentation of the reward cue, and there is only a weak non-significant cross-temporal generalisation between the first and second delay. A dynamic, rather than stable, representation of space returns around the time of the go cue. In the low time constant population, coding is always dynamic, so no stable state is established. Dotted lines encircling areas of strong coding indicate significant cross-temporal stability (p<0.05, **Methods**).

The resulting pattern of generalisation can distinguish between different working memory models, as indicated by the exemplars in **Fig4a**. The first example shows a ‘stable attractor’ model on SR trials^7^. Soon after the spatial cue is visible, a stable state of network activity forms specific to each spatial location. This pattern of activity generalises (i.e., the “off-diagonal” regions of the matrix) throughout the time the stimulus remains in working memory (illustrated by red colour from stimulus presentation onwards). A revised ‘stable subspace’ version of this model incorporates a dynamic component during the cue period, with a stable state only present from the delay period (Exemplar 2)^13^. In this version, spatial coding during cue presentation doesn’t generalise to later periods in the trial, but a stable attractor is formed around the time of stimulus offset. A third exemplar shows what may happen if this stable subspace were to be disrupted by the presentation of the reward cue (’distractible subspace’). A final example shows a purely ‘dynamic coding’ model ^38,46^, whereby dynamic on-diagonal selectivity maintains an active representation of spatial information across time, but this never reaches a fixed point of stable network activity (i.e., lack of off-diagonal shading).

The pattern produced by the activity of the VLPFC high tau subpopulation exhibited elements consistent with both stable and dynamic coding (**Fig 4b, d**)^13,48^. Coding from the spatial cue period was not positively correlated with the subsequent delay, consistent with dynamic activity during the initial encoding phase. Surprisingly, neural activity was anti-correlated between the cue and the delay (largest cluster, p<0.0001; cluster based permutation test), suggesting the way the network encodes spatial information reverses between cue presentation and delay. This selectivity pattern reversal was also evident in VLPFC reward coding, but was not present in any other PFC area despite strong reward selectivity in ACC and OFC (**Supplementary Fig1**).

Despite this dramatic reversal of selectivity from the cue to delay periods, a stable state of cross-temporal generalisation was established in the high tau subpopulation during the first delay epoch which was sustained through the reward cue epoch (**Fig4b**; largest cluster, p<0.0001; cluster based permutation test). This finding is consistent with the VLPFC high tau subpopulation demonstrating attractor-like working memory activity in classical tasks without intervening stimuli^1,7,13^. However, the cross-generalisation of maintained spatial information was disrupted during the subsequent reward delay epoch on SR trials, and there was no significant generalisation between the activity in the first and second delay (**Fig4b**, no candidate clusters). The fact that network activity in the VLPFC high tau subpopulation is dynamic at cue presentation, then exhibits a reversed stable state of generalisation which is disrupted following the distractor (reward cue), suggests VLPFC network activity is not performing the function of a conventional attractor for spatial working memory^7^.

Compared with high tau cells, the VLPFC low tau subpopulation had much more transient dynamics (**Fig4c, e**). Although there is weak on-diagonal selectivity, this does not extend off the diagonal, consistent with dynamic coding. The spatial selectivity in the high tau subpopulation was significantly more stable over time during the post-stimulus delay and shortly after (largest clusters, SR p = 0.0002, RS p = 0.0135; cluster based permutation test; **Supplementary Fig2**). In summary, of all of the subpopulations across the PFC areas we examined, only the high tau VLPFC subpopulation formed a stable spatial mnemonic representation, but the additional task element of a salient distractor allowed us to show that this state was inconsistent with current attractor models.

### Anti-correlation between Cue and Delay Period Activity

Recent work has suggested that stable population activity can co-exist alongside strong temporal dynamics during the initial encoding phase^13^. This can occur if the mnemonic representation is established at the time of the cue but is accompanied by a transient, orthogonal pattern of activity. These results would appear inconsistent with the reversal of spatial coding we observed in the VLPFC high tau population between cue presentation and delay. To examine this issue in more detail, we correlated activity within the VLPFC high tau subpopulation across time within each condition (**Fig5a**, see **Methods**)^13^. This showed a strong positive correlation across the whole trial, including between cue and delay periods (asterisk on **Fig5a**). This suggests that within a given spatial location, VLPFC high tau firing rates were stable and correlated across the trial (as opposed to the instability and reversal of mnemonic coding across the trial evident in **Fig4**). Whilst this may be taken as evidence against a reversal of selectivity patterns, we reasoned this positive correlation may be largely driven by the intrinsic firing rates of the neurons (e.g. a neuron which is high firing during the cue may also be higher firing during the delay even if it is modulated across the trial). By demeaning activity across conditions for each neuron and repeating the analysis, we revealed an anticorrelation in the activity of high tau VLPFC neurons between the spatial cue and delay periods (asterisk on **Fig5b**, see Methods). The high cross-trial correlations observed in **Fig5a** are therefore likely driven by neurons possessing relatively consistent firing across the trial.

**Figure 5:**
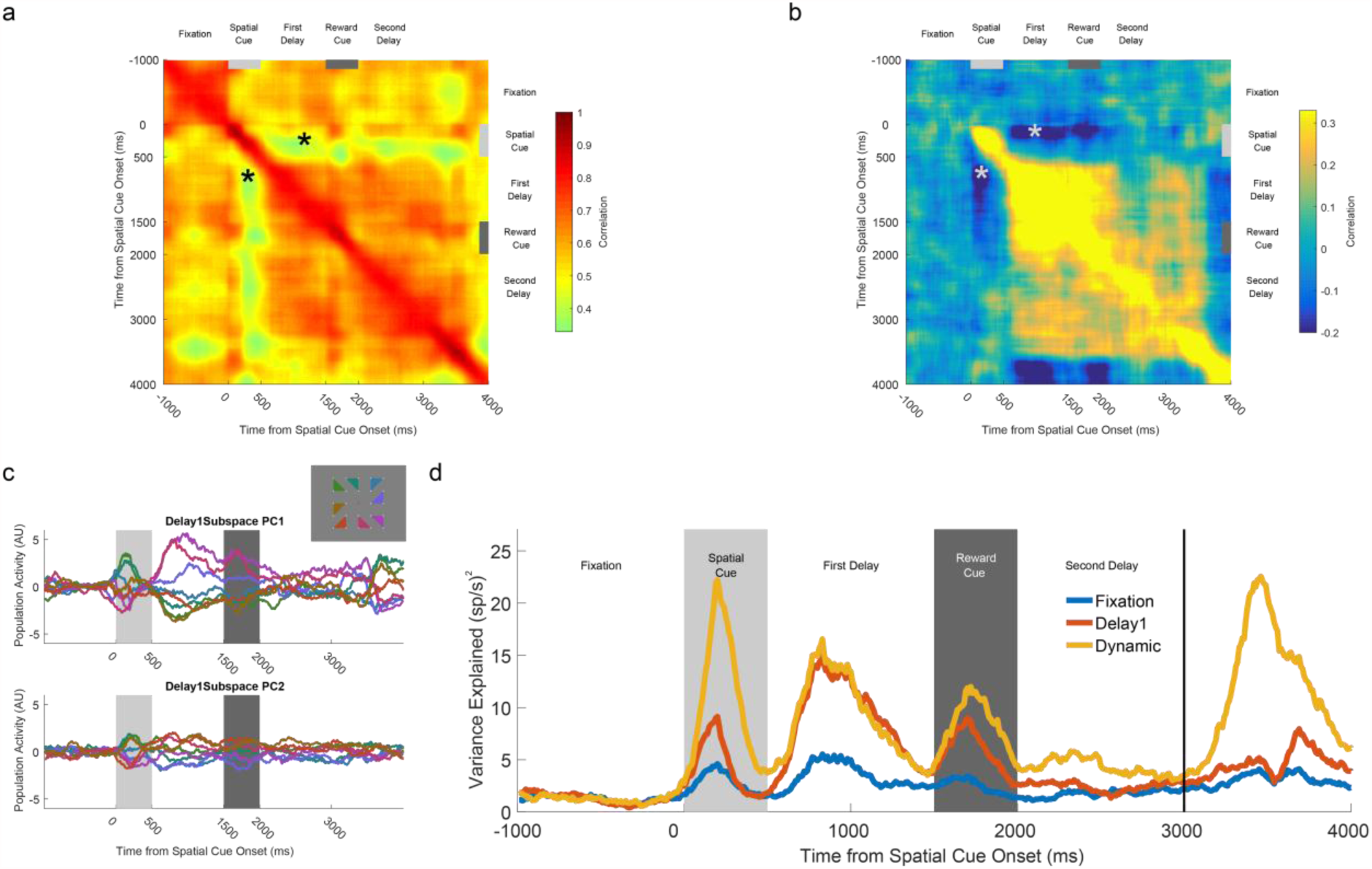
VLPFC high time constant population reverses its spatial coding between cue presentation and the subsequent delay. **a)** Within-condition correlation of neural firing across time for SR trials. All bins are positively correlated with each other, suggesting neural firing is stable across time. Note positive correlation between cue period and delay (asterisk). **b)** Within-condition correlation analysis where activity for each neuron was demeaned across each of the spatial locations. There now exists a negative correlation between the time of the spatial cue presentation and the first delay (asterisk). **c)** Reversal of VLPFC high time constant spatial tuning between cue and delay. A mnemonic subspace was defined by time-averaged delay one activity. The across-trial firing for each condition was projected back onto the first and second principal axes of this subspace. While the conditions remain well-separated on both principal axes during the first delay, the subspace does not generalise well into the second delay as activity from the different conditions converges. At the time of the cue, the conditions appear separable, but in the reverse configuration from that during the delay. The inset shows the geometric location of each spatial location that appeared on the screen. **d)** The stimulus variance captured by three different subspaces is displayed. The fixation subspace is defined by time-averaged activity in the 1000ms before cue presentation. This should represent a chance-level amount of variance explained. The Delay1 subspace is defined by time-averaged activity from 500ms to 1500ms after cue presentation. The dynamic subspace is defined separately at each individual time point. The dynamic subspace explains a much greater amount of variance during the cue period, illustrating that there is little consistency in the activity patterns between cue and delay epochs. However, the Delay1 subspace captures as much variance as the dynamic subspace during the first delay, suggesting the VLPFC high tau population activity has settled to a stable state by this point.

To further examine the stability and pattern of spatial selectivity across the trial using an alternate method, we employed principal component analysis (PCA). Previously, this method revealed a mnemonic subspace that was stable from cue onset through the delay period^13^. The mnemonic subspace was defined by time-averaging delay period activity for each stimulus condition for each neuron and running PCA across conditions (conditions × neurons matrix). Projecting data from the cue period into this subspace still enabled decoding of spatial position, supporting the proposal that the stable activity in the delay period is already established during cue presentation^13^.

We tried to replicate this PCA approach in the high tau VLPFC subpopulation (**Fig5c-d**, see Methods), by defining the subspace based upon time-averaged delay one activity in the SR trials. We then projected neural firing from across the trial onto the first two principal axes (**Fig5c**). If the mnemonic representation is stable, all traces should be fairly fixed and separable across time (as in ref^13^ FigS3). During the first delay, there is a stable representation of mnemonic information, as all conditions are separable within this subspace. The representation of space is also shown to be geometrically consistent with the spatial environment, with the activity for nearby spatial locations clustered in the subspace. However, supporting our previous analyses, projecting neural activity from the cue period into the subspace didn’t lead to a reliable spatial code. Remarkably the spatial conditions were separable in the cue period, but in the opposite direction to that observed during the delay period. This pattern was also replicated for reward coding on RS trials, suggesting this reversal is a general pattern of VLPFC coding between cue and delay periods, and not limited to spatial selectivity (**Supplementary Fig3**). To quantify the reliability of the SR Delay 1 subspace, we calculated the variance explained by projecting data at each timepoint (**Fig5d**). Unlike previous findings^13^, the mnemonic subspace in the delay explains only a small proportion of variance during the cue period.

In short, we find little evidence that the VLPFC high tau subpopulation forms a stable subspace maintaining information from cue onset through the delay. Rather, the population geometry reverses its selectivity pattern for both reward and spatial information between the cue and delay periods (**Fig4b, 4d, Fig5c-d, Supplementary Fig3**), before forming a stable subspace that maintains WM-related information across the initial delay before the subsequent cue (distractor) period.

### Cross-Task Generalisation

Thus far we have demonstrated that only high tau VLPFC neurons exhibit stable *cross-temporal generalisation* of mnemonic information. We next explored whether there was *cross-task generalisation* between SR and RS trials. Previous studies have demonstrated task-specific PFC activity to identical stimuli when they cue a different response^49,50^. However, whether the pattern and stability of population activity depends on the order in which identical information (cueing the same response) is received remains unknown. To explore this, we used data from one trial type as a training set, and data from the other trial type as a test set. This analysis allowed us to test, for example, whether the population pattern for spatial selectivity that emerges in delay one of SR trials (**Fig4b**) is similar to the population pattern for spatial selectivity in delay two of RS trials. This analysis also allowed us to test whether the population pattern in delay two was similar across both trial types; at this point in the trial, the subjects have processed the same information and are required to prepare the same response.

**Fig6a** depicts three possible exemplars for cross-task generalisation. As demonstrated in **Fig2**, VLPFC has spatial coding on both trial types, thus if there is “no across-task generalisation” this would mean there are multiple network patterns of spatial selectivity capable of supporting correct performance. In “stimulus/delay-locked cross-task generalisation”, the population pattern in the spatial cue and subsequent delay periods is similar across trial types. In this scenario, spatial location could be readout identically across trial types using activity post-stimulus presentation (red colour on heatmap), but because spatial selectivity on SR trials is disrupted by the reward cue (**Fig4**), distinct readout weights would be required at the time of response. In “action-dependent cross-task generalisation”, the population selectivity pattern is similar across trial types only during delay two and the saccade response. This may occur if a different route through neural state space is taken on the two trial types, but the routes converge and the same common endpoint is reached by delay two.

**Figure 6:**
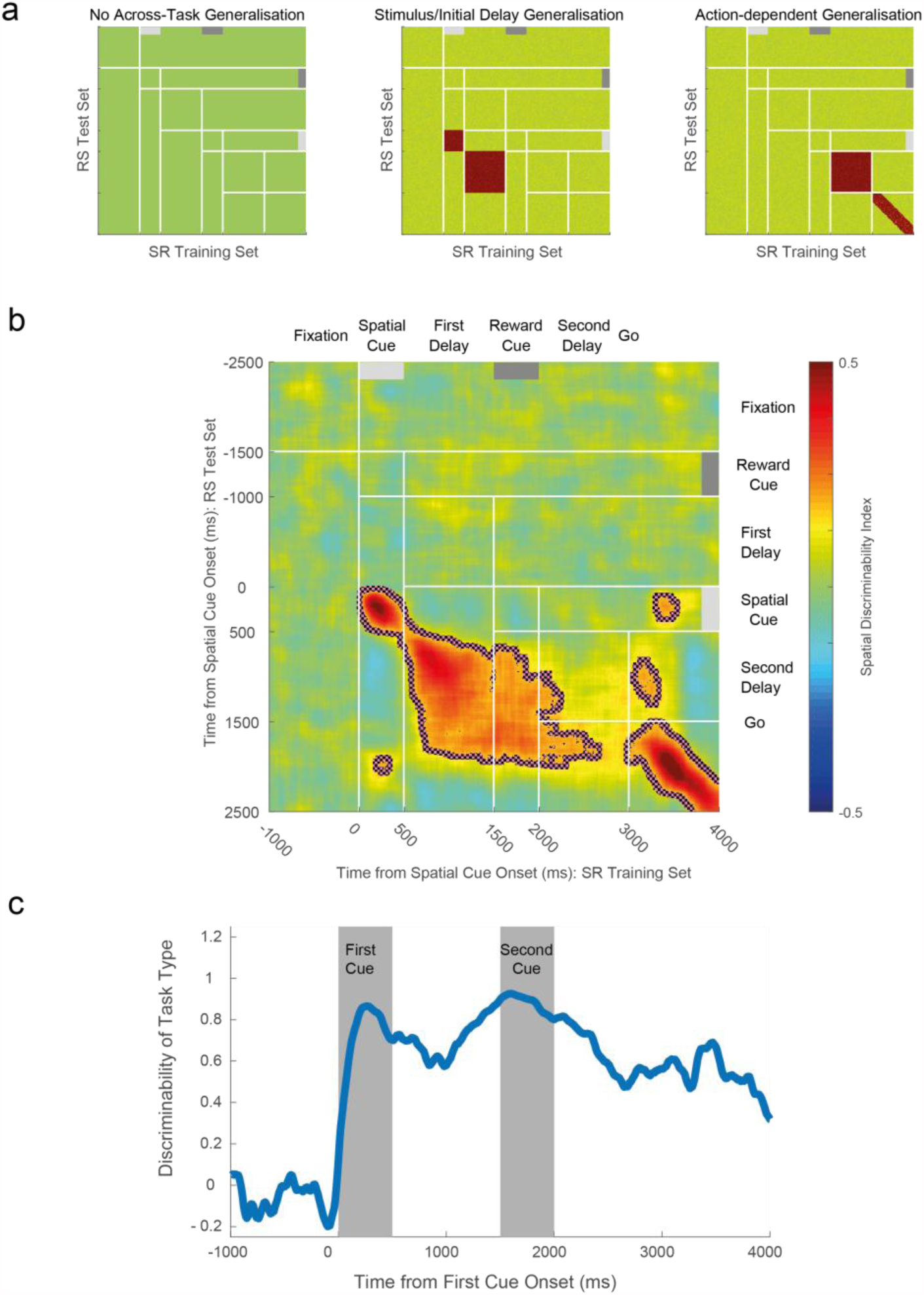
Cross-generalizability of working memory across trial types. By using data from SR trials as a training set for a classifier, and data from RS trials as a test set, the generalizability of spatial coding across task types can be studied. **a)** Exemplars of how population activity may generalise across trial types. If there is no across-task generalisation, spatial position cannot be decoded from neural activity recorded on the other trial type. If there is stimulus-locked generalisation, spatial position can be decoded by activity from the other trial type; however, it is relative to cue presentation so the decoding is displaced off of the diagonal. If there is action-dependent generalisation, neural activity generalises along the diagonal in the second delay and response epochs as subjects prepare and execute their saccade. **b)** Cross-generalizability in VLPFC is primarily locked to the presentation of the stimulus. Spatial position cannot be decoded from activity during the second delay period, implying distinct population codes on the two trial types in the delay immediately prior to response. Only once the action is initiated (at the go cue), does a cross-trial generalisation appear on the diagonal. Dashed lines encircling areas of strong coding indicate a significant cross-generalizable stability (p<0.05, see Materials and methods). **c)** Decoding task type. The task the subjects are performing can be accurately decoded from VLPFC neural activity, throughout the trial. This is particularly important during the second delay, as at this point the subject has been exposed to the same visual stimuli, just in reverse order.

We performed this analysis on all recorded VLPFC neurons. The activity pattern of this population was primarily consistent with stimulus locked generalisation (**Fig6b**). This is because there is strong cross-task generalisation between when the spatial cue is presented and during the initial one-second mnemonic period following that (Cue period p<0.0001; Delay period p<0.0001; cluster based permutation tests). There is then little cross-task generalisation in delay two, indicating distinct activity patterns in this epoch between the two tasks. We confirmed a strong representation of trial type during delay two using a separate decoding algorithm, which discriminated activity related to trial type (**Fig6c**, see **Methods**). These results indicate that a different set of read-out weights for working memory of spatial location would be required from VLPFC activity for correct performance on the two trial types, implying multiple, independent task-specific neural states can support working memory.

#### Single neurons switch between reward and spatial selectivity

Thus far, the results suggest a heterogeneous and primarily dynamic account of working memory activity within the PFC population. To examine the underlying pattern of this population heterogeneity, we analysed single neuron selectivity for different task features. This analysis explored how strong and sustained WM selectivity patterns were in individual neurons^8,48^, how these WM representations were affected by the presentation of a second salient cue (which may induce selectivity competition), and whether neural activity in delay two encoded a combination of task variables^25,26^.

To quantify single-neuron encoding of both reward and spatial information, we ran a separate oneway Kruskal-wallis test for space and reward at each timepoint (**Fig7a, b**). Encoding at each timepoint was determined significant through a cluster-based permutation test (see **Methods;** cluster-forming threshold, p <0.05). This allowed us to plot whether each neuron was encoding space, reward or both factors at any given point in time (**Fig7c, d**). On the SR trials, a large proportion of VLPFC neurons were selective for spatial location during cue presentation or the subsequent delay (**Fig7a**, top). These neurons had heterogeneous onset latencies and most were transiently selective, as opposed to sustaining a spatial representation across time. Strikingly, many of these spatially selective neurons subsequently coded reward size later in the trial (**Fig7a**, bottom). This is consistent with the VLPFC population analysis (**Fig2**) showing that the most recently presented stimulus is encoded, as opposed to the task-relevant spatial information necessary for correct performance. A similar result was also observed on RS trials, where many reward-selective neurons (**Fig7b**, top) subsequently encoded spatial location (**Fig7b**, bottom). This suggests that the population-level patterns we observed (**Fig2**) arise because single PFC neurons are involved in multiple distinct cognitive functions^25^, as opposed to different subpopulations representing different task-related factors becoming active at different stages of the trial.

**Figure 7:**
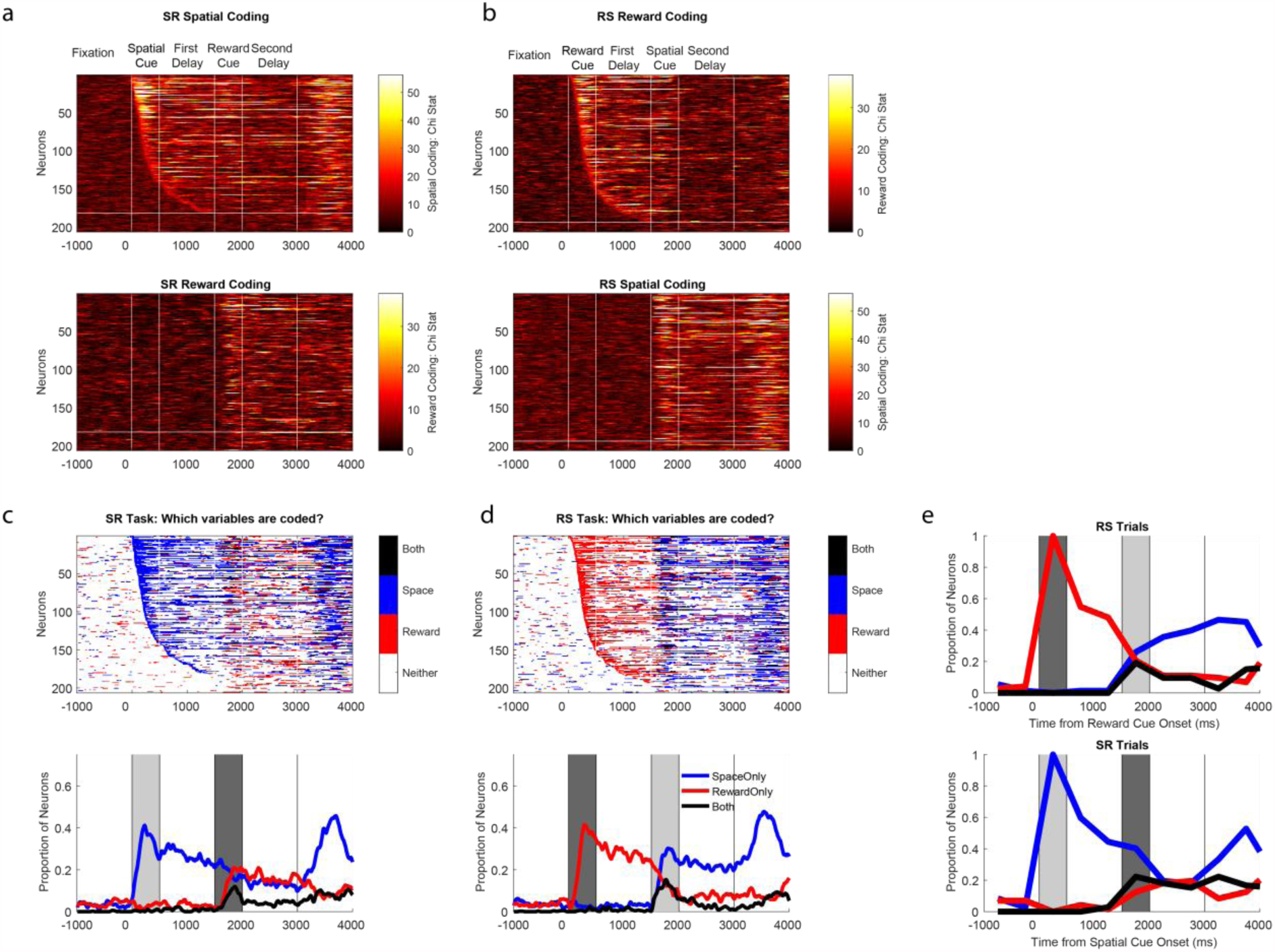
Flexibility of single-neuron selectivity. **a**) SR Trials: Single neuron coding. The top plot shows the spatial coding of individual ventrolateral PFC neurons; each row of the matrix represents single neuron selectivity. Neurons are sorted by their latency for spatial encoding; all neurons above the horizontal white line were selective for space either during cue presentation or the first delay. For many of these cells, selectivity is transient; few code space across extended periods of the trial. Furthermore, a large proportion of these neurons subsequently become selective for reward at cue two/delay two (neurons are sorted in the same order in the panel below). **b**) RS Trials: Single neuron coding. Neurons are now sorted by their latency for reward encoding, with all neurons above the white line selective during cue presentation or the first delay. The top panel shows reward encoding, which again is primarily transitory in nature. The bottom panel shows that many of the neurons initially coding reward go on to code the spatial-location when this cue is presented. Fraction of neurons selective for either or both task factors across SR (**c**) or RS (**d**) trials. Presentation of the second stimulus reduces the number of neurons selective for the initially presented cue. **e**) Switching of selectivity across a trial. Neurons are included in this analysis if they were selective during the presentation of the first cue. The selectivity pattern of these neurons is profiled across time. On SR trials, only a minority of cue selective neurons retain an exclusive representation of space across the entire trial; many neurons gain reward coding, some at the expense of spatial selectivity, and others in addition to this.

The ability of PFC neurons to encode both reward and spatial information may highlight neuronal flexibility, or the facility to code multiple factors concurrently. **Figs7c-d** characterise the proportion of neurons simultaneously coding spatial and reward information. During the presentation of the second cue, some neurons appeared to multiplex reward and spatial information. To establish the nature of this mixed selectivity, we ran a 2-way ANOVA to explore any interaction effects (**Fig8**, see **Methods**). It could be that neurons code both factors with a non-linear interaction^21,25^, exhibiting a different pattern or degree of spatial coding at each reward level. Alternatively, both factors could be coded simultaneously without an interaction^51^ (e.g. similar pattern of spatial selectivity for each reward level resting upon a different baseline firing rate). We found little evidence for non-linear mixed selectivity. Instead, there was a positive correlation between selectivity for space and reward at the time of the second cue (**Fig8**), implying most neurons that exhibit mixed selectivity for multiple factors do so as a linear combination.

**Figure 8.**
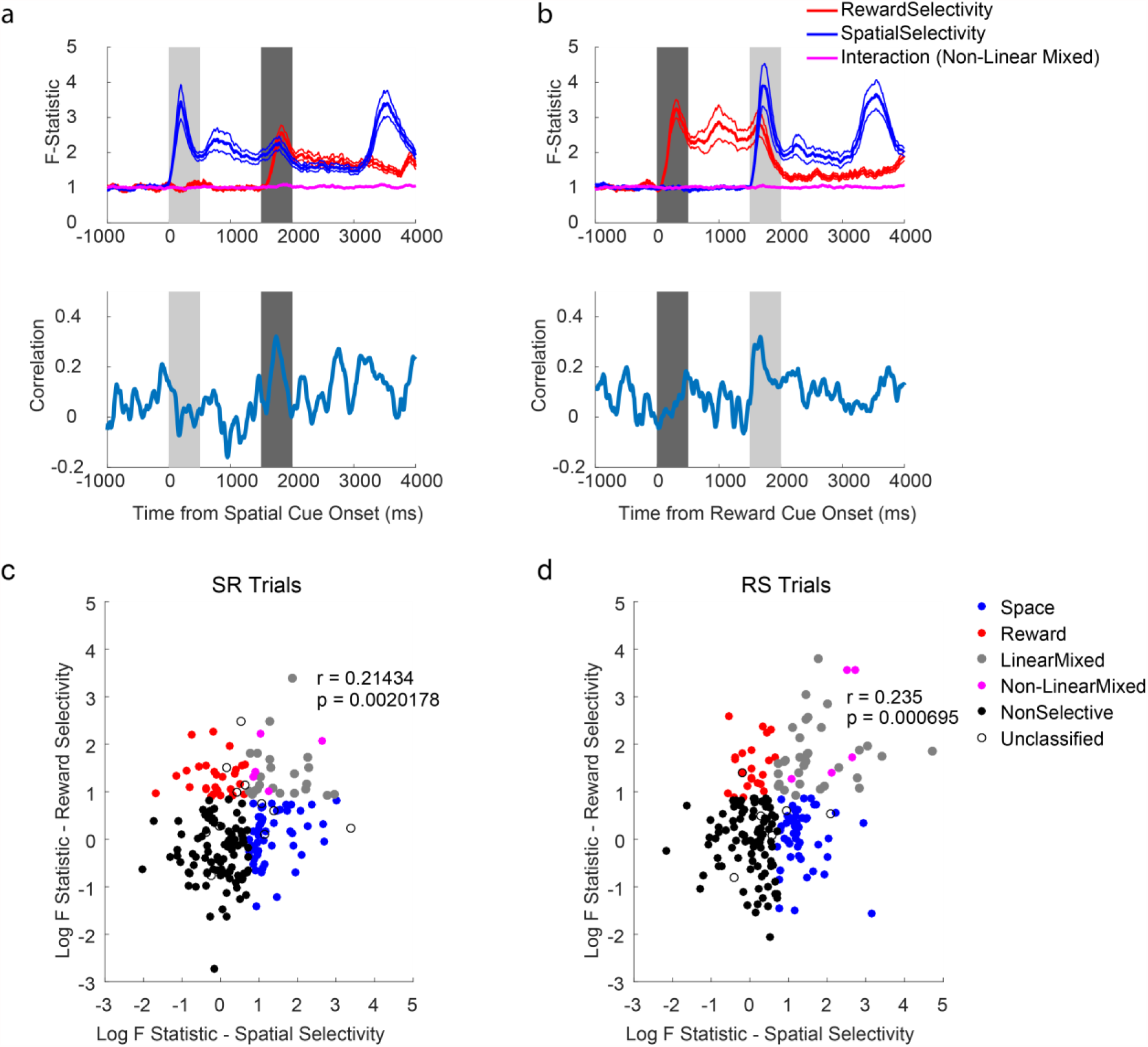
Absence of non-linear interactions between reward and spatial selectivity on SR (a) and RS (b) trials. The mean population F-statistics from a sliding 2-way ANOVA with an interaction term are plotted (±SEM; top panel). The interaction term between both factors does not change from that during pre-trial fixation. However, when there is a positive correlation between reward and spatial selectivity F-statistics at the time of Cue 2 (Bottom panel), indicating linear mixed selectivity. Spearman correlation of Spatial and Reward coding F-Statistics during Cue 2 for **c**) SR trials; **d**) RS trials. To complement the above analysis, we performed a single spearman correlation between the raw F-statistics from a 2-way ANOVA of spike-counts during the second cue. Each dot represents a neuron appropriately coloured. Space and Reward neurons were required to have only one significant main effect (p<0.05) and a non-significant interaction (p>0.05). Linear mixed neurons were required to have two significant main effects (p<0.05) and a non-significant interaction (p>0.05). Non-linear mixed neurons were required to have two significant main effects (p<0.05) and a significant interaction (p<0.05). Non-selective neurons had no significant effects (p>0.05). A very small proportion of neurons did not meet these criteria so were unclassified. F-statistics have been log-transformed for illustrative purposes.

This flexibility of single-neuron selectivity patterns appears inconsistent with more traditional accounts of PFC function during working memory. To quantify the proportion of neurons exhibiting stimulus-specific selectivity throughout the trial, we split the data into eight 500ms epochs from fixation onset until the response was cued. We ran a separate Kruskal-wallis test on the average firing rate of each neuron across these epochs. A subpopulation of neurons with selective responses during the initial cue presentation was defined (n=73 for reward, n= 72 for space). The proportion of this subpopulation selective for each factor was then calculated for all other epochs (**Fig7e**). On SR trials, this showed that only 18.06% of the spatially selective neurons at cue one are selective for spatial location alone by the end of the second delay. Around the same proportion (15.28%) had additionally gained a representation of reward size, whilst a further 19.44% had no significant spatial selectivity and switched to coding reward. The majority (47.22%) of spatially selective neurons at cue one were non-selective by the end of delay two. Thus unlike classical notions of working memory being supported by sustained selectivity^1,2^, our results suggest single neurons do not maintain sustained working memory correlates^9^, at least not in cases where other task-relevant or salient information may compete for neuronal representation.

## Discussion

Here we used a spatial working memory task with a distracting reward cue to test whether working memory (WM) is subserved by persistent activity in single neurons or by dynamic activity across a neural population. This task design with a distractor allowed us to specifically contrast these different hypotheses of WM coding. A recent cortical attractor model of WM would suggest a dynamic cue-related response followed by a stable state of fixed network activity specific to the mnemonic stimulus^13^. This model would predict that if changes in this stable state were induced by distractor presentation, this would compromise the WM representation. This constraint does not apply to WM models that do not rely on stable network states. Of the four PFC subregions examined, mnemonic selectivity during the delays was present only in VLPFC neurons, and this was present only in the subpopulation of neurons with high time constants. Within these VLPFC neurons, the pattern of both reward and spatial selectivity reversed from the cue to delay epochs, where it then became stable and generalised across time. However, once the reward cue was presented, spatial selectivity was largely quenched and instead the VLPFC population switched to coding the salient reward information. These results demonstrate that high tau VLPFC neurons are capable of stable selectivity that could serve WM functions, but that in contexts where multiple behaviourally relevant stimuli are available, VLPFC neurons flexibly code the focus of current attention^26,37,39^.

Both attractor^13,19,52^ and synaptic models^14^ of working memory stress the importance of a recurrent network architecture. By using the decay of autocorrelation of spiking activity during a fixation period as an unbiased metric of intrinsic persistent activity, we demonstrate that neurons with higher time constants (taus) are more likely to exhibit working memory-related selectivity, but only in the VLPFC population. The VLPFC high tau subpopulation had stable selectivity during the initial delay period following stimulus offset, whereas the low tau subpopulation exhibited dynamic coding. Importantly, any distinction between the high and low tau VLPFC subpopulations was only evident during this mnemonic phase, ruling out the possibility that high tau cells are simply more task-selective. These results build upon recent work showing PFC neurons with higher taus have a greater role in decision-making and the maintenance of reward information over extended time periods^33^, highlighting a broader role for high time constant neurons subserving extended cognitive processes. These findings would therefore appear supportive of theories proposing that cortical attractor networks fulfil WM functions^7,13^.

However, we also observed several features of the data which suggest VLPFC activity is incompatible with current attractor models. Firstly, we showed that VLPFC reverses both its spatial and reward tuning between cue presentation and the subsequent delay. Previous studies have shown that cue and delay dynamics are distinct^13,38,48^, but our discovery that the tuning geometry reverses between cue and delay appears novel. This finding is also inconsistent with a stable subspace spanning both cue presentation and memory^13^. The inverted tuning geometries may reflect a mechanism to dissociate stimuli currently in the environment and those held within memory^53^, or alternatively a mechanism to load information into working memory from an initial state of dynamic sensory encoding.

By probing the effect of a salient reward cue on the stability of mnemonic representations, we were able to further test whether cortical attractors in PFC provide a mechanism for distractor-resistant WM. It was shown that the intervening reward cue quenched the WM selectivity pattern in the VLPFC population. A recent report similarly showed that a task-irrelevant distractor morphed spatial selectivity of PFC neurons^21^; however this irrelevant distracting cue could be instantly dismissed and was not encoded. The PFC population activity, although morphed with respect to activity pre-distraction, could therefore continue to maintain a strong mnemonic representation. In our paradigm, the reward cue acted as a more ethologically-valid distractor with behavioural relevance. Reward anticipation is known to activate a large proportion of neurons in prefrontal cortex^22,43,54-59^, and many neurons holding the spatial representation flexibly switched to code the reward. This suggests that different neural mechanisms may be required to maintain WM when a distracting stimulus also carries behavioural relevance and activates neurons across PFC. This WM mechanism seemingly eludes current attractor models, which predict distractor-resistant spatial selectivity.

The dynamic switch of VLPFC activity to coding the behaviourally relevant distractor provides further evidence that PFC neurons can be tuned to multiple diverse cognitive factors and that they can flip between them within the course of a trial^25,27,48,60^. It also suggests previous studies concluding PFC neurons are resistant to distraction do not generalise to more behaviourally salient stimuli^28-30^. Here we use a reward-predictive cue presented at the fixation spot, as opposed to a peripherally flashed target^29^ or stimulus^21,28^ which is irrelevant to the task. The flexibility with which VLPFC neurons changed the factor they encoded also has implications for accounts of mixed selectivity^21,25,51^. Shortly after the second stimulus was shown, there was evidence for neurons encoding a combination of factors. However, we found the majority of this mixed selectivity was linear^51^, as opposed to non-linear^21,25^.

Inverted tuning between cue and delay, a weakening of a stable mnemonic representation by a distracting cue, and neurons flexibly encoding both factors all suggest VLPFC activity is incompatible with existing cortical attractor models^13^. There are several possible interpretations of the WM activity we observed across PFC. Although WM-related activity and WM-deficits following brain damage are both most commonly associated with LPFC^4,5,61^, it is conceivable that classical distractor resistant stable activity was present in a PFC region we did not sample. However, we sampled a large expanse of LPFC including both banks of the principal sulcus (PS: areas 9/46d, 946v), and several millimetres of cortex both dorsal (area 9) and ventral (areas 45a, and 47/12) to PS, as well as parts of the medial (ACC) and ventral (OFC) PFC. Mnemonic activity has been observed in other brain regions, such as the parietal cortex^62,63^. However, this activity is more sensitive to distraction^29,64^, and parietal inactivation produces comparatively modest WM deficits relative to PFC, suggesting it plays less of a role in WM processes^6,65,66^. A further possibility is that we may have missed stable, persistent activity in PFC because of a more anterior recording location than previous studies^41^. We also consider this interpretation unlikely. Recent studies recording more posteriorly in LPFC including the frontal eye field have shown that selectivity for a remembered spatial location is not stable when either multiple mnemonic stimuli are subsequently presented or a distractor appears^21,67^. Instead, we note that the vast majority of tasks reporting stable coding do so during a delay period where there is only one mnemonic representation to be maintained and no intervening stimuli^2,3,13,48^. Had our study similarly terminated at the end of delay one on SR trials (**Fig4b**), our findings would be highly consistent with findings of these tasks^13^. Crucially, without presentation of a distractor stimulus, both the most recently presented stimulus and the posited locus of the subject’s attention are confounded with working memory^39^. Our findings suggest stable mnemonic representations are present in PFC, specifically in high tau VLPFC neurons, but that these neurons can also flexibly switch which information they encode as other behaviourally relevant variables compete for the subject’s attention.

Of the PFC regions studied, VLPFC mnemonic representations were the strongest, and the only ones present during the second delay of SR trials, although in an altered state relative to the initial delay. The question therefore remains how WM is achieved on this task. One possibility, although not directly verifiable with our data, is that a PFC region maintains a representation of the mnemonic stimulus in an activity silent state^14,15^. Alternatively, PFC may be essential for setting up a stable mnemonic spatial representation during the initial delay which can then be transmitted to oculomotor regions to prepare a saccade, akin to activity for reaching movements^68^. Either way, our data are incompatible with PFC maintaining WM in cortical attractor networks throughout a delay interrupted with a behaviourally relevant distractor. This provides novel neurophysiological evidence that stable activity states within PFC may be more tightly associated with the most-recently presented behaviourally-relevant stimulus, rather than the contents of working memory.

## Methods

### Subjects and neurophysiological procedures

Neurophysiological procedures and task design have been reported previously^36,37^. In brief, two male rhesus macaques (Macaca mulatta) served as subjects. Single neuron recordings were taken from four regions of prefrontal cortex (**Fig1b**) including dorsolateral prefrontal cortex (DLPFC, n=209), ventrolateral prefrontal cortex (VLPFC, n=206), orbitofrontal cortex (OFC, n=152) and the anterior cingulate cortex (ACC, n=198). Histological reconstructions of the precise locations of all recorded neurons have been reported previously^36,37^. We randomly sampled neurons and did not attempt to pre-select neurons based on responsiveness to enable a fair comparison of neuronal properties between different brain regions.

### Task

A detailed overview of the task structure has been described elsewhere^36,37^. We monitored eye position and pupil dilation during the task using an infrared system (ISCAN). Subjects first fixated a central cue for 1000ms before two cues were presented sequentially, each for 500ms, each followed by a 1000ms delay. One of the cues was a spatial location that the subject had to hold in working memory, and the other indicated how much reward the subject would receive for correct performance of the trial. We used 24 different spatial positions and two different reward-predictive picture sets, each cue indicating one of five reward levels (**Fig1a**). The 24 spatial targets were regularly distributed in a 5 x 5 matrix centred at the fixation spot, with each position separated by 4.5° of visual angle. The positions were collapsed into eight locations forming triangles to allow for sufficient trials for the decoding analyses. On Space-Reward (SR) trials, the spatial position was shown first followed by the reward cue, whereas on Reward-Space (RS) trials the cues were presented in the reverse order. If subjects maintained fixation through both of the cue and delay periods, the fixation cue changed colour and the subject could initiate a saccade to the remembered spatial location (**Fig1a**). If the saccade terminated within 3° of the remembered target and was maintained in this location for 150ms, a reward was delivered and the trial was recorded correct. Trials where fixation was maintained but the saccade failed to terminate in the remembered location were recorded as errors. Different trial types and conditions were randomly intermingled. Subjects completed ~600 correct trials per day.

### Data-analysis

Single-neuron activity during a 1000ms fixation period was used to assign time constants (**Fig1c-d**)^33^. Single unit responses were time-locked to the onset of the fixation period of successfully completed trials. Fixation-period rasters were divided into 20 discrete, successive 50ms bins. The spike count for each neuron within each bin was calculated for each trial. Pearson’s correlation coefficient was used to compute the across-trial correlation of spike-counts between all of the bins. For each single neuron, this produced an exponential decay when autocorrelation was plotted as a function of time lag between bins (as in **Fig1d**). The decay of the autocorrelation was fitted to the data using the following equation^34^:

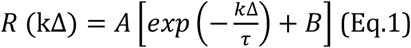

In which kΔ refers to the time lag between time bins (50 to 950ms) and τ is the time constant of the neuron (**Fig1c**), when data from one autocorrelogram is fitted, or the cortical area when data from all neurons within that area are fitted together (**Fig1d**). Neurons from all areas, particularly ACC, showed evidence of lower correlation values at the shortest time lag^33^. This may reflect refractoriness or negative adaptation^34^. To overcome this, fitting started from the largest reduction in autocorrelation (between two consecutive time bins) onwards.

All recorded neurons were included in the population-level time constant analysis (**Fig1d**). Single neurons were assigned a time constant if their autocorrelogram could be reasonably described by an exponential decay^33^. Neurons were therefore automatically excluded if they had a fixation firing rate of <1Hz or no decline in their autocorrelation function in the first 250ms of time lags (28 of 765 excluded). Neurons were also excluded if the fitting produced extreme parameters (A > 1.2, A < 0, τ>1000, τ<10; 156 of 737 excluded). Finally, this was followed by a process of visual inspection where a further set of neurons were excluded which were considered to possess autocorrelation functions poorly characterised by an exponential decay (172 of 581 excluded). This left 141 DLPFC, 157 VLPFC, 73 OFC and 38 ACC neurons for analysis. Two independent observers completed this process, blind to each neuron’s functional properties and recording location. The majority of excluded cells were recorded in ACC, where many neurons’ autocorrelation functions were flat, possibly reflecting a timescale longer than could be indexed with a 1-seond foreperiod. In VLPFC, which is the brain region where most analyses were performed, only 23.8% of all recorded neurons were excluded. All results were replicated without the visual inspection exclusion criteria.

A multivariate decoding approach was used to investigate population-dynamics of working memory coding^38^. Decoding was performed separately for different task-types (i.e. SR or RS) and different task features (i.e. space and reward). For each neuron, correct trials were split equally into a training set and a test set. Within each set, trials were grouped according to the relevant feature to be decoded (either eight spatial groups or five reward levels). Neuronal firing rate for each of these conditions *(Conds)* was averaged across trials for each neuron producing a vector length **Conds**. The pairwise difference between neural firing in each of the conditions was calculated. For eight spatial locations (five reward levels) this produced 28 (10) pairwise differences *(PWDs)*. The Pearson’s correlation coefficient for each *PWD* was calculated across neurons between the training set and the test set. These correlation coefficients were averaged using Fisher’s Z-transformation to produce a single correlation-coefficient quantifying either reward discriminability or spatial discriminability. This process was repeated for each timepoint, so that the temporal profile of decodability could be evaluated (**Fig2-3**). A similar analysis was used to probe if the task being performed could be decoded (**Fig6c**).

Cluster-based permutation tests were used to correct for multiple comparisons while assessing the significance of time-series data^33,69^. Discriminability metrics were compared between the high and low tau subpopulations using Fishers-Z transformation (**Fig3**). This yielded a test-statistic at each timepoint. Test statistics were divided into ten, non-overlapping 500ms epochs beginning at fixation onset. Consecutive bins in each analysis window with an uncorrected (cluster-forming) threshold of p<0.05 (one-tailed) were defined as candidate clusters. The size of the clusters were compared to a null distribution constructed using a permutation test. Neurons assigned to each subpopulation were randomly permuted 10,000 times and the cluster analysis was repeated for each permutation. The length of the longest cluster for each permutation was entered into the null distribution. The true cluster size was significant at the p<0.05 (p<0.01) level corrected if the true cluster length exceeded the 95^th^ (99th) percentile of the null distribution. A cluster’s significance was determined to be p<0.0001 if its length exceeded all those in the null distribution. A similar method was used to compare discriminability to chance levels (**Fig2**). Consecutive bins in each analysis window with an uncorrected (cluster-forming) threshold of p<0.01 (two-tailed) were defined as candidate clusters. In this case, permuted clusters were calculated by shuffling the order of neurons in each of the PWDs in the test set.

The multivariate decoding approach allowed us to also probe the cross-temporal stability of mnemonic representations (**Fig4**). The discriminability measure described above involved correlating the PWDs calculated at the same timepoint for a training and a test set. In the cross-temporal analysis, a ***timepoints × timepoints*** matrix was constructed where the training set at each timepoint was tested at all other timepoints^33,38^. In **Fig4** the matrix of correlation coefficients was averaged across the diagonal in order for the data to reflect both training-to-test and test-to-training trial projections. To probe the stability of population coding in this analysis, cluster-based permutation tests were used. Neighbouring pixels in each analysis window with an uncorrected (cluster-forming) threshold of p<0.01 (two-tailed) were defined as candidate clusters. The null distribution was generated by the same permutation method as in **Fig2**. To compare the stability between high and low time constants, a two-dimensional version of the Fishers-Z transformation method described above was used (**Supplementary Fig2**).

Independent to selectivity measures, neural firing rate was correlated across the trial (**Fig5a, b**). Firing rate for each condition (eight spatial locations, five reward levels) was correlated across neurons between each timepoint pair. A separate training and test set were defined based upon a split half of the trials. The matrix of correlation coefficients plotted represents the average (using Fisher’s Z-transform) value across all of the conditions (**Fig5a**). For **Fig5b**, prior to performing the correlation, neural firing rate was demeaned within each condition and timepoint for each neuron.

Principal component analysis (PCA) was used to perform a state space analysis (**Fig5, Supplementary Fig3**)^13^. Each subspace was defined using a training set of data averaged across half of the available trials for each neuron and tested using data from the remaining half. This makes stimulus-variance captured non-arbitrary (**Fig5d**) and explains why only a minimal amount of variance is explained in fixation before stimulus presentation. For each neuron, firing rate on training set trials was averaged for each condition for each timepoint. For the fixation and delay one subspaces, activity was averaged across the relevant timepoints (Fixation: −1000 to 0ms relative to cue onset; Delay One: 500ms to 1500ms relative to cue onset). This produced a *Conds × Neurons* matrix. Activity was demeaned across conditions for each neuron. PCA was then performed over conditions to define a low-dimensional coding subspace for the two epochs within a high-dimensional neural state space. For the dynamic subspace, firing was not averaged across timepoints and the PCA was performed separately at each timepoint. Therefore, a slightly different subspace is produced for each time point. Once the principal components have been defined, we projected the left-out test set data onto the principal axes of the subspaces (**Fig5c**). The plotted traces therefore display a lowdimensional representation of the trajectory of population activity in the subspace across time.

To assess the generalizability of the delay one subspace, we plotted the stimulus variance (**SV**) it captured across the trial relative to the fixation and dynamic subspaces (**Fig5d**). SV was calculated using the following formula:

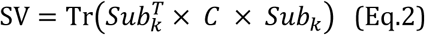

In which **Sub_k_** refers to the subspace defined from training data (limited to the first k principal axes) and **C** refers to the across-stimuli covariance matrix of the test data. In our analyses, we used one fewer principal axes than the number of conditions (Space: k =7; Reward: k = 4).

For the preliminary single-neuron encoding analyses (**Fig7a-d**), a one-way kruskal-wallis test was performed for spatial location and reward size at each time point. A cluster-based permutation test was performed to test for significance (**Fig7c-d**). Consecutive bins in each analysis window with an uncorrected (cluster-forming) threshold of p<0.05 were defined as candidate clusters. In this case, permuted clusters were calculated by shuffling the relevant feature (spatial location or reward size) across trials. To probe whether neurons coding for both factors simultaneously demonstrated either linear or non-linear mixed selectivity, we performed a two-way ANOVA (**Fig8**).

Several graphs with time series data were smoothed across time bins for illustrative purposes (**Fig 2; Fig3; Fig6c; Fig7c-d**, bottom half). A moving average spanning five 10ms bins was used. However, all statistical tests were performed on the unsmoothed data.

## Acknowledgements

SEC was supported by the Middlesex Hospital Medical School General Charitable Trust. JDW was supported by funding from NIMH R01-MH097990 and NIDA R21-DA035209. LTH was supported by a Sir Henry Wellcome Fellowship from the Wellcome Trust (098830/Z/12/Z). SWK was supported by NIMH (F32MH081521) and by a Wellcome Trust New Investigator Award (096689/Z/11/Z). We thank S Mark for comments on an earlier draft of this manuscript.

---

**Supplementary Figure 1:**
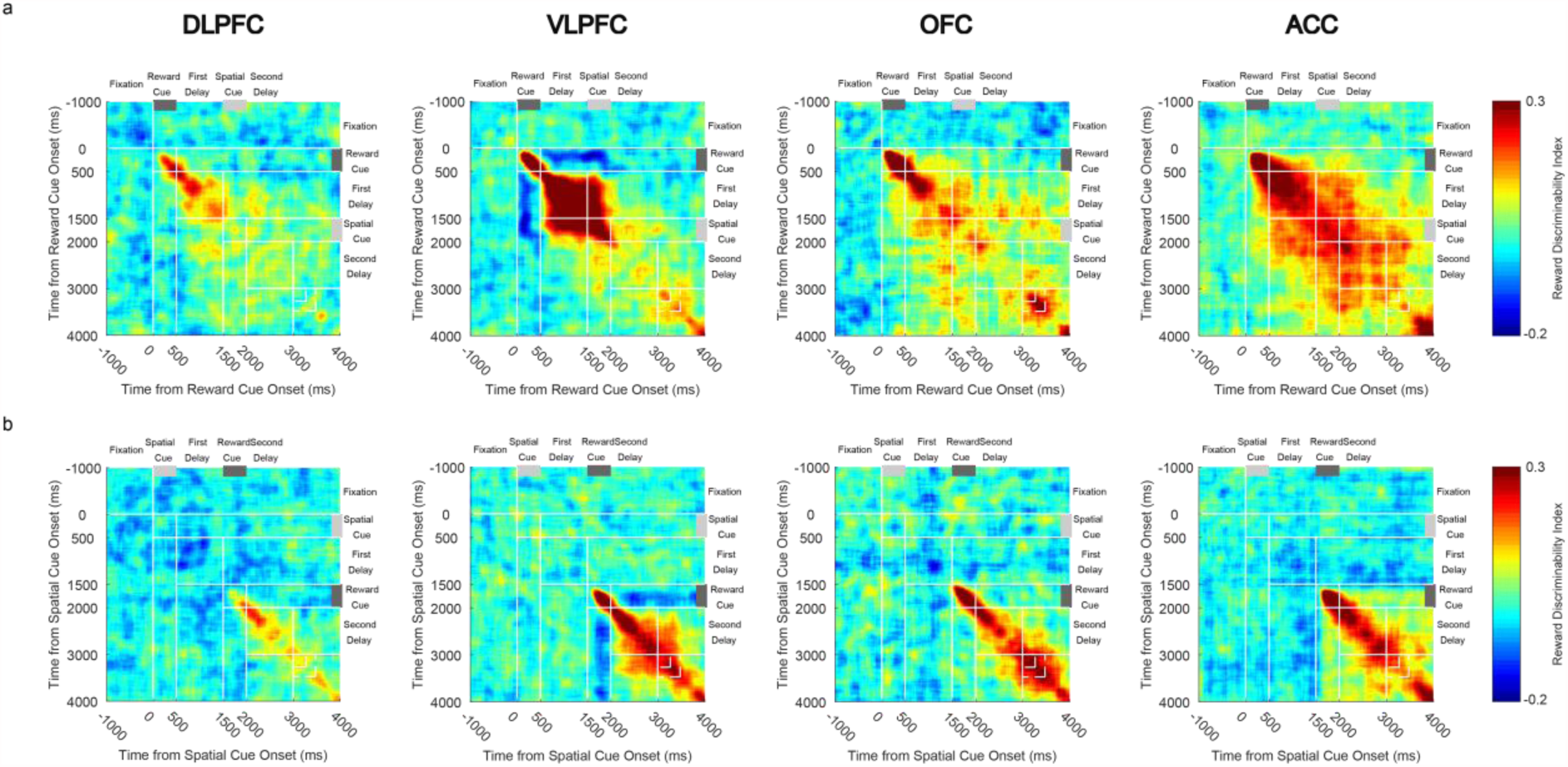
Cross-temporal dynamics of reward selectivity by brain region and task (a, RS task; b, SR task). All brain areas studied have neural activity representing reward size. Only VLPFC shows a reversal of reward tuning between the cue epoch and the subsequent delay. This feature of coding is present on both trial types.

**Supplementary Figure 2:**
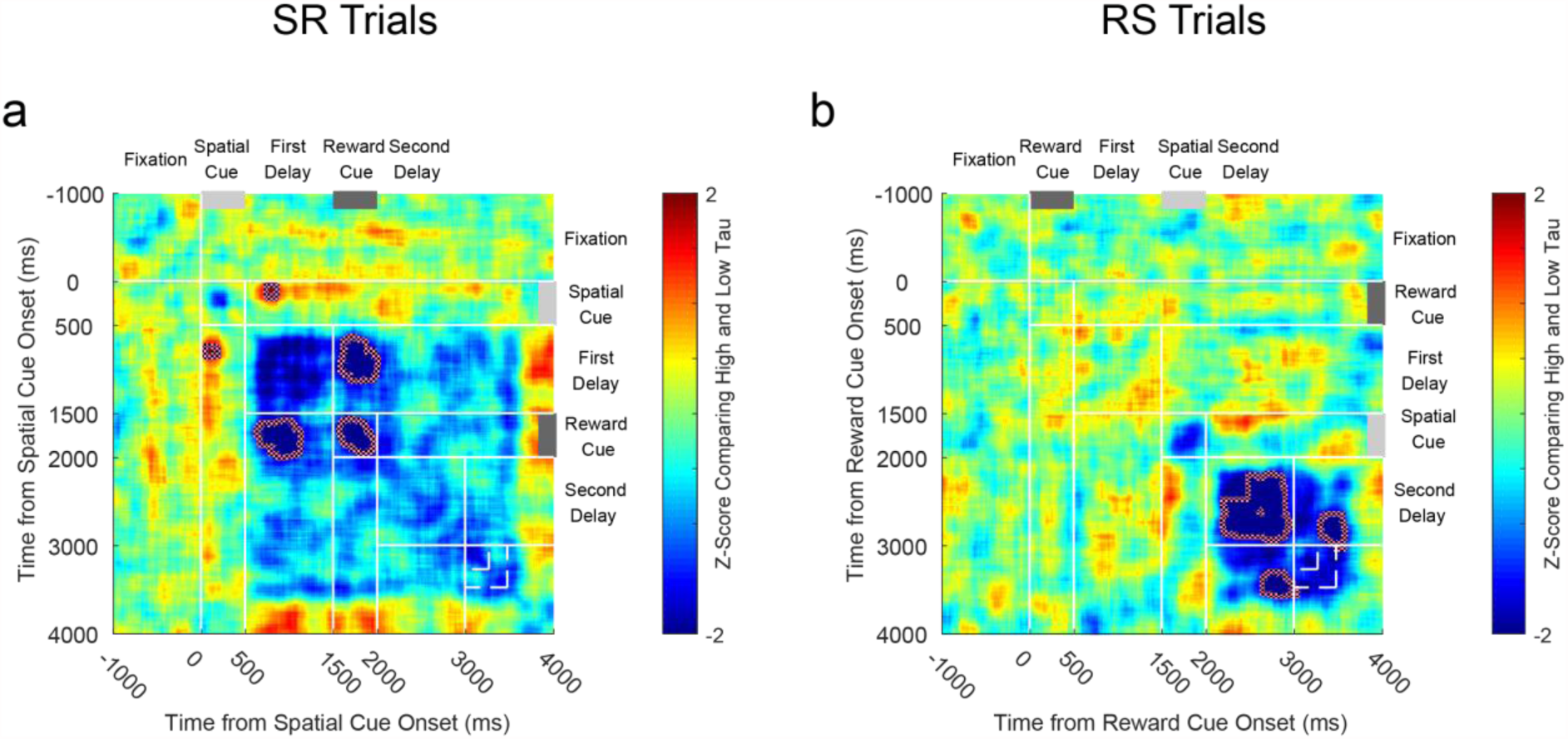
Comparison of VLPFC High and Low tau cross-temporal spatial coding for a) SR trials; b) RS trials. Negative z-scores illustrate stronger coding in the high tau population. Coding of spatial location is more stable for the high tau population between the first delay and the reward cue of SR trials (largest cluster, p = 0.0002; cluster based permutation test, see **Methods**), and during the reward cue of SR trials (largest cluster, p = 0.0086; cluster based permutation test). There is a stronger switch in coding between the spatial cue and the first delay in high tau cells (largest cluster, p = 0.0079; cluster based permutation test). On RS trials, there is a more stable coding in high tau cells during the second delay (largest cluster, p = 0.0135), as well as between this time and the reward onset (largest cluster, p = 0.0120; cluster based permutation test). Dotted lines encircling areas of strong dissimilarities in coding indicate a significant difference in cross-temporal stability between high tau and low tau populations (p<0.05, see Materials and methods).

**Supplementary Figure 3:**
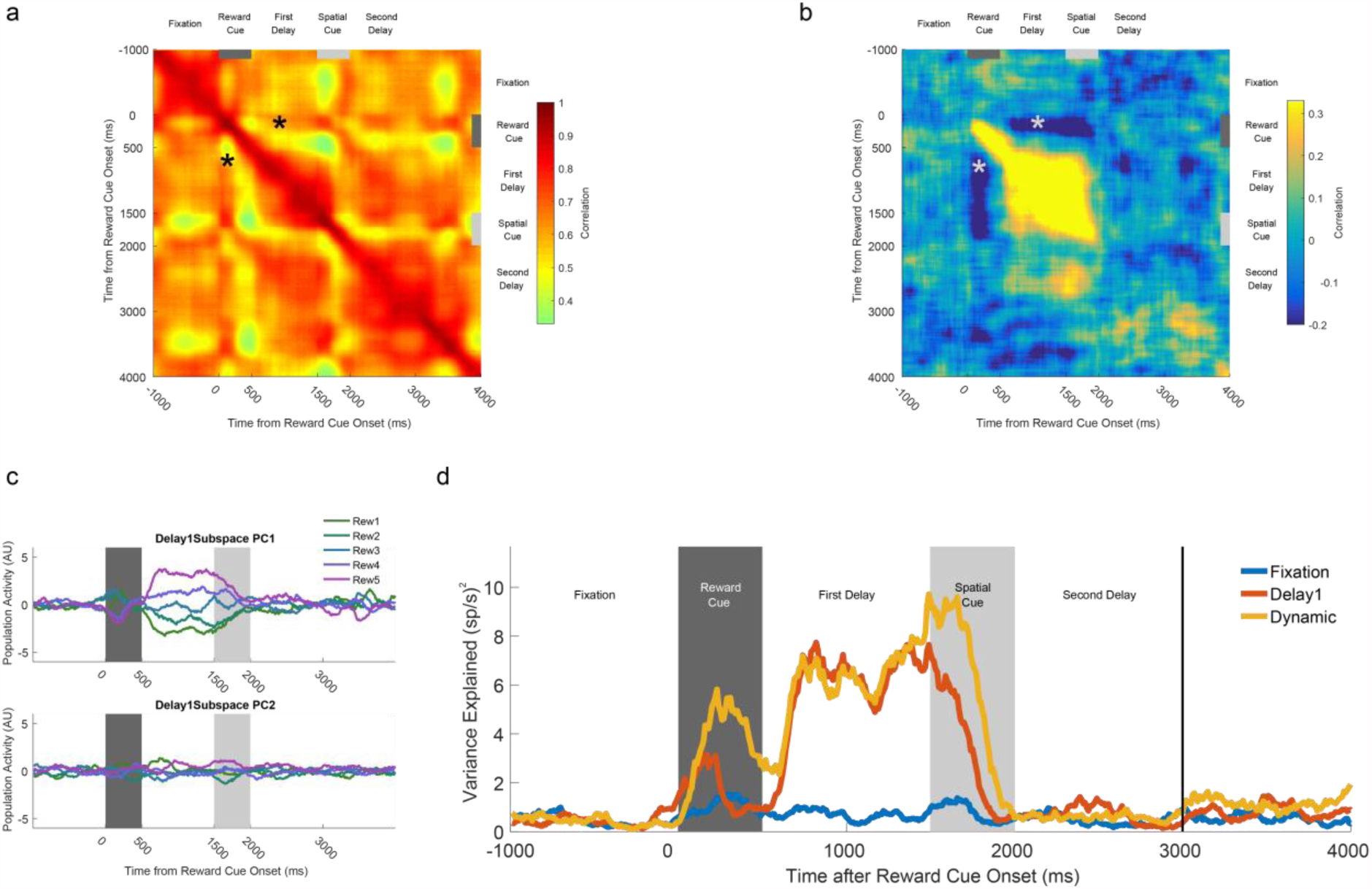
VLPFC high time constant population reverses its reward coding between cue presentation and the subsequent delay. **a)** Within-condition correlation of neural firing across time for RS trials. All bins are positively correlated with each other, suggesting neural firing is stable across time. Note positive correlation between cue period and delay (asterisk). **b)** Within-condition correlation analysis where activity for each neuron was demeaned across each of the reward sizes. There now exists a negative correlation between the time of the reward cue presentation and the first delay (asterisk). **c)** Reversal of VLPFC high time constant reward tuning between cue and delay. A mnemonic subspace was defined by time-averaged delay one activity. The across-trial firing for each condition was projected back onto the first and second principal axes of this subspace. While the conditions remain well-separated on the first principal axis during the first delay, the subspace does not generalise well into the second delay as activity from the different conditions converges. At the time of the cue, the conditions appear separable, but in the reverse configuration from that during the delay. **d)** The stimulus variance captured by three different subspaces is displayed. The fixation subspace is defined by time-averaged activity in the 1000ms before cue presentation. This should represent a chance-level amount of variance explained. The Delay1 subspace is defined by time-averaged activity from 500ms to 1500ms after cue presentation. The dynamic subspace is defined separately at each individual time point. The dynamic subspace explains a much greater amount of variance during the cue period, illustrating that there is little consistency in the activity patterns between cue and delay epochs. However, the Delay1 subspace captures as much variance as the dynamic subspace during the first delay, suggesting the VLPFC high tau population activity has settled to a stable code by this point.

